# Lipid osmosis, membrane tension, and other mechanochemical driving forces of lipid flow

**DOI:** 10.1101/2024.01.08.574656

**Authors:** Yongli Zhang, Chenxiang Lin

## Abstract

Nonvesicular lipid transport among different membranes or membrane domains plays crucial roles in lipid homeostasis and organelle biogenesis. However, the forces that drive such lipid transport are not well understood. We propose that lipids tend to flow towards the membrane area with a higher membrane protein density in a process termed *lipid osmosis*. This process lowers the membrane tension in the area, resulting in a membrane tension difference called *osmotic membrane tension*. We examine the thermodynamic basis and experimental evidence of lipid osmosis and osmotic membrane tension. We predict that lipid osmosis can drive bulk lipid flows between different membrane regions through lipid transfer proteins, scramblases, or other similar barriers that selectively pass lipids but not membrane proteins. We also speculate on the biological functions of lipid osmosis. Finally, we explore other driving forces for lipid transfer and describe potential methods and systems to further test our theory.

## Lipid transfer through lipid transfer proteins and scramblases

Lipid trafficking is essential for the formation and proper function of organelles in the cell [1]. Lipids are primarily synthesized in the endoplasmic reticulum (ER) and transported to various organelle membranes. Damaged membranes or organelles are delivered to and degraded in lysosomes. Such lipid trafficking can be vesicle-dependent or -independent. While the former has been well understood, which also mediates the trafficking of proteins and other molecules, the latter has just been emerging as a key pathway specific for lipid transfer [1-3]. The nonvesicular lipid transport is mediated by a large, conserved family of lipid transfer proteins (LTPs) [4]. They transfer lipids at the membrane contact sites where two membranes are brought into 10–30 nm by LTPs and other membrane tethering proteins [5]. LTPs are categorized into bridge and shuttle LTPs based on their structures, functions, and mechanisms of lipid transfer (Figures 1a & 1b). Bridge LTPs (BLTPs) contain taco-like grooves of 10–30 nm in length lined with hydrophobic residues [6,7] (Figures 1a & 1c). With both ends attached to membranes, they act as a conduit to allow lipids of different species to flow quickly through the groove. Bridge LTPs not only help regulate lipid compositions but also mediate organelle biogenesis and dynamics associated with large changes in membrane areas and morphologies [8] (Figure 1d). Shuttle LTPs contain hydrophobic cavities that bind one or a few lipids at a time (Figure 1b). They are generally tethered to one membrane at the membrane contact sites and transfer lipids between the two membranes, often by sequentially exchanging lipids at both membranes, thereby regulating lipid compositions [9-11] (Figure 1e). However, both kinds of LTPs transfer lipids in the cytosolic membrane leaflets. Lipid scramblases equilibrate lipids between two leaflets and may cooperate with bridge LTPs to transfer lipids [12-15] (Figures 1a & 1f). The scramblases provide an intramembrane hydrophilic channel or thin the membrane for lipids to flip between two leaflets [16-18]. Whereas the effects of lipid transfer are widely observed, it remains unclear what drives the directional lipid flow and how such lipid flow alters the lipid compositions of membranes [1,3]. Some lipid transfer processes are coupled with ATPases that pump lipids against their concentration gradients, such as MsbA, lipopolysaccharide transport (Lpt) proteins, and lipid flippases [17,19]. However, LTP-mediated lipid transfer can be decoupled from ATP or other energy sources. For example, BLTPs can transfer lipids between inner and outer membranes of bacteria when ATP is depleted in the cells [20,21]. Instead, the directional lipid transfer likely results from the continuous or discontinuous flow of lipids along gradients of their chemical potentials across different membranes or membrane domains, which is our focus in this work. Therefore, in the following, we will not distinguish lipid transfer and lipid flow.

**Figure 1.**
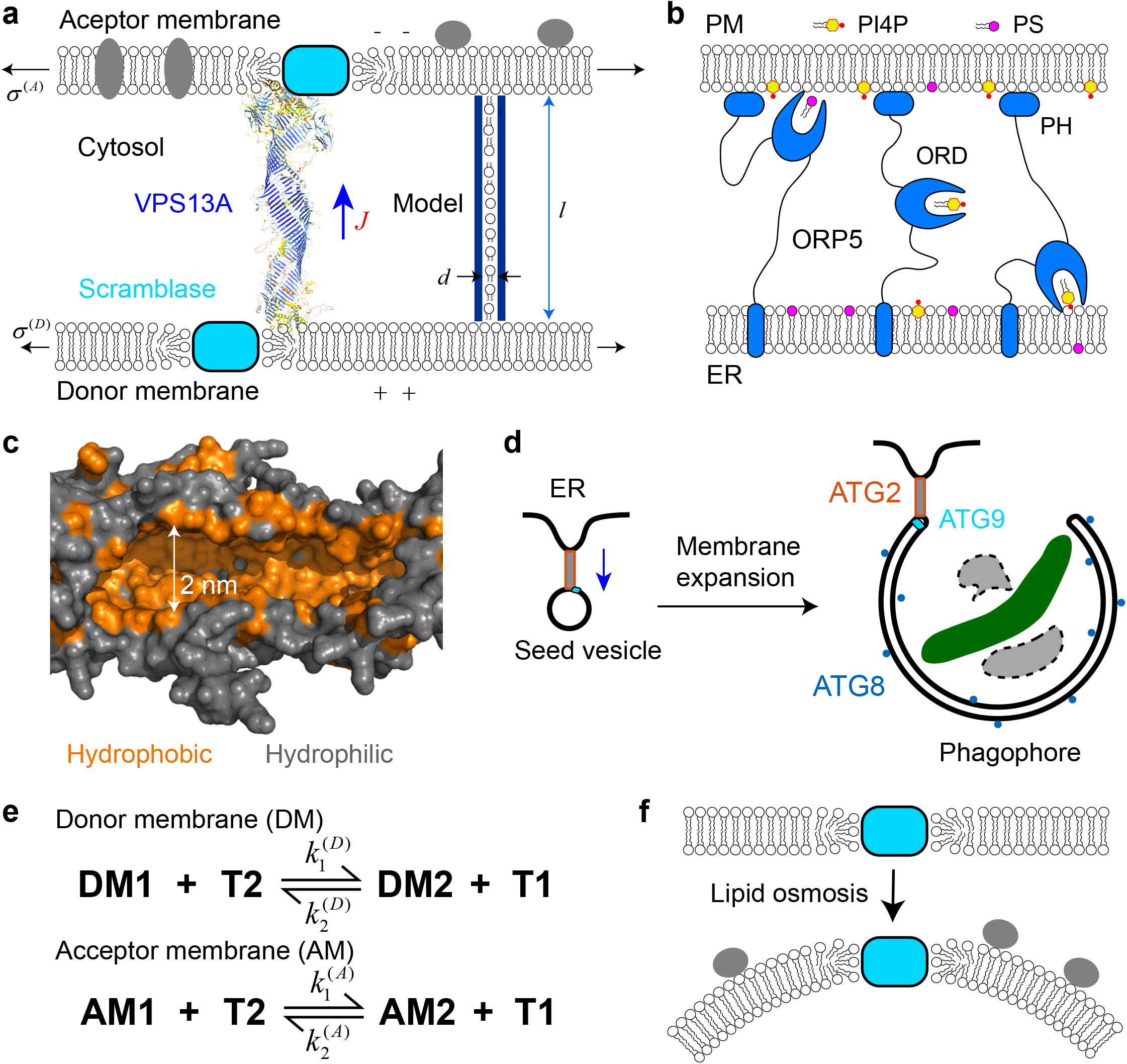
Lipid transfer proteins (LTPs) and scramblases transfer lipids between different membrane leaflets. **(a)** A bridge LTP (BLTP), shown here by the AlphFold structure of a representative BLTP VPS13A [56], acts as a lipid conduit to allow bulk lipid flow from one membrane to the other. The flow (with its direction indicated by the blue arrow) may be driven by the gradient of membrane tension, electric potential, or membrane protein density as indicated. The channel on the right represents an idealized BLTP used to estimate its lipid flow rate. **(b)** A representative shuttle LTP ORP5 transfers phospholipids PS and PI4P between the plasma membrane (PM) and the ER through two lipid-exchange processes. ORP5 contains an N-terminal pleckstrin homology (PH) domain that binds to the PM, a middle OSBP-related domain (ORP) that binds to PS or PI4P, and a C-terminal transmembrane domain in the ER. Three snapshots of the lipid transfer are shown. **(c)** A segment of the hydrophobic groove of the bacterial BLTP YhdP predicted by AlphaFold. **(d)** During macroautophagy, BLTP ATG2 and the scramblase ATG9 mediate fast lipid flow from the ER to the double-membraned phagophore for its membrane expansion, which eventually engulfs damaged organelle (green) or protein aggregations (gray) for their degradation. The enzyme-catalyzed lipidation of ATG8 proteins (blue dots) on the phagophore membrane is required for the membrane expansion. **(e)** The two lipid-exchange processes are characterized by two sets of rate constants, which may be equal or different, depending upon the membrane environments. DMi, AMi, and Ti with i=1 or 2 denote the i-th lipid species in the donor membrane, acceptor membrane, and shuttle LTP, respectively. **(f)** Lipid osmosis can occur between different leaflets of the same membrane mediated by scramblases, which may cause membrane bending.

### Lateral lipid flow between different membrane domains through protein fences

Lipids also flow laterally between different membrane domains, which may be driven by new forces in addition to the well-known membrane tension gradient. Cell membranes contain transient or permanent domains with distinct sizes and compositions of lipids and proteins, which are often separated by barriers formed by protein complexes [22]. These barriers border the membrane domains and block or hinder the diffusion of membrane proteins across the barriers but generally allow faster passage of lipids [23,24]. For example, the plasma membrane of a neuron is highly compartmentalized by distinct diffusion barriers (Figure 2a). These barriers are located at the initial segment between the cell body and the axon and the neck of the dendritic spine [25,26]. In addition, the axon membrane is supported by actin rings evenly distributed along the entire length of the axon, which also hinders the diffusion of membrane proteins [26,27]. Similarly, septin rings help compartmentalize plasma membranes at the bases of cilia and the cleavage furrow during cell division [28-30], while tight junctions maintain apical and basolateral membranes in epicedial cells [22]. Intracellular cytoskeleton or extracellular matrix attaches plasma membranes to form dynamic fence-like structures [31,32]. These fences permit lipids to flow in and out but confine the mobile proteins for a significant time before they escape the fences, creating highly heterogeneous protein distributions in plasma membranes [33]. Other barriers separate vesicle precursors from the plasma membrane during endocytosis, lipid droplets from the ER membrane, the ER exit site from the ER membrane, mitochondrial cristae from the inner boundary membrane (Figure 2b-e) [34,35], and the nuclear envelope from ER [36]. Many peripheral proteins are recruited to these isolated membrane domains to facilitate organelle biogenesis [37,38]. These barriers show common features with LTPs in that they selectively pass lipids between two membranes or membrane domains with different densities of membrane proteins.

**Figure 2.**
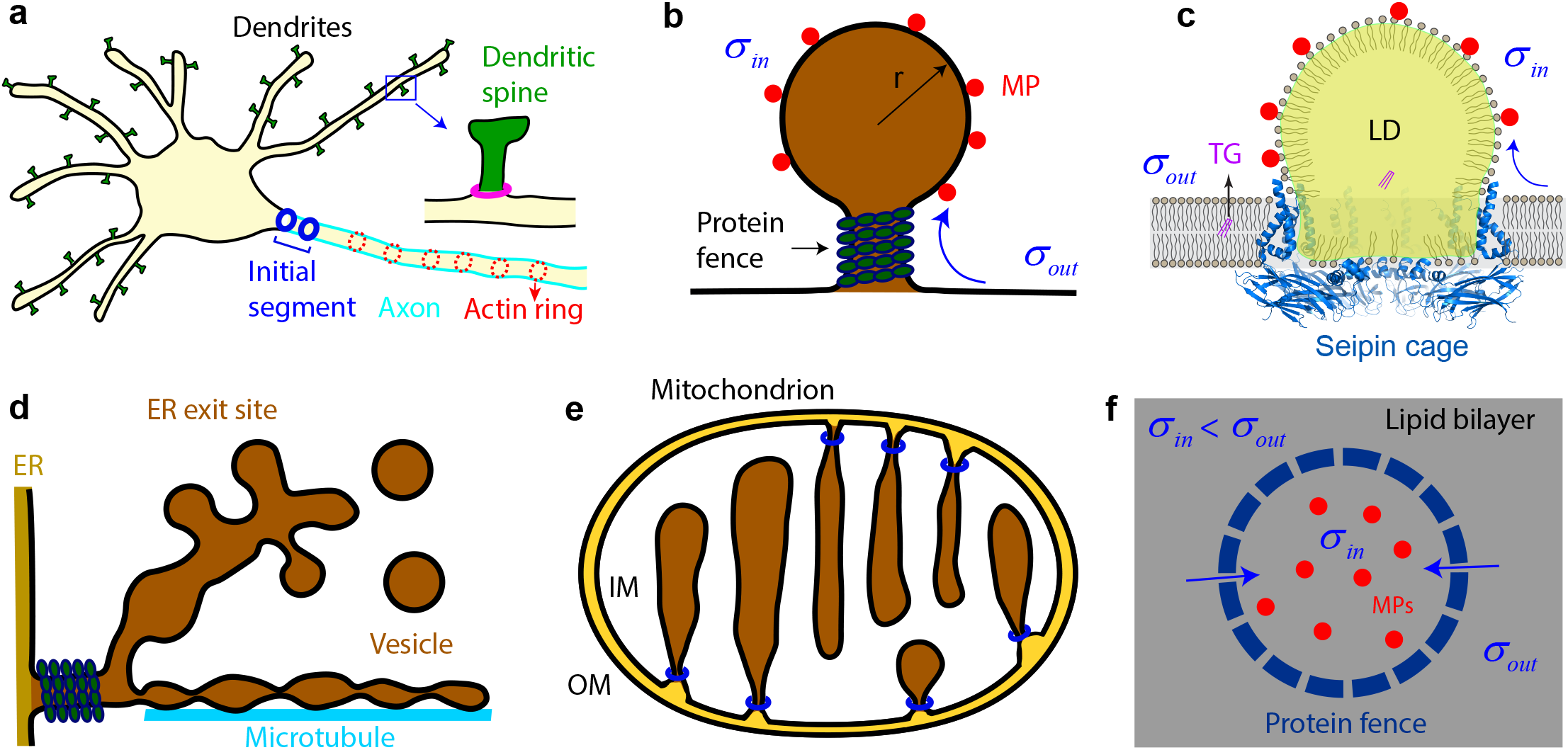
Ubiquitous protein barriers separate cell membranes into domains with distinct functions and membrane protein densities and compositions. **(a)** The axon and dendritic spines are separated from the neural cell body at the initial segment and dendritic spine neck, respectively, by physical barriers (blue and magenta circles) that limit protein diffusion across the membrane domains. **(b)** The vesicular precursor crowded with membrane proteins (red circles) is connected to the plasma membrane through a protein-coated neck. Lipid osmosis (blue arrow) may facilitate the vesicular membrane expansion. **(c)** Seipin forms a decameric fence-like structure in the ER membrane to mediate lipid droplet (LD) formation between the two ER leaflets. Each droplet contains a neutral lipid core (highlighted in yellow) containing triglycerides (TG) covered by a phospholipid monolayer and various membrane proteins (red dots). **(d)** The ER exit site is a specialized ER domain that serves as a hub for membrane trafficking. The domain is connected to the ER membrane by a neck coated by proteins including COPII and TANGO1. **(e)** The mitochondrial inner membrane invaginates to form cristae with protein-coated crista junctions (marked by blue circles) connected to the inner boundary membrane. **(f)** Lipid osmosis occurs through a protein fence that selectively passes lipids but not membrane proteins (blue dashed ring), leading to lower equilibrium tension of the membrane inside the fence with enriched proteins.

Membrane tension gradient has long been recognized as a major driving force for lateral lipid flow within the same membrane [23,24,32,39] and is expected to also serve as a driving force for the lipid flow across different membranes through LTPs or leaflets through scramblases (Figures 1a & 1f). However, the membrane tension of some plasma membranes varies significantly in different areas tested without significant lipid flow between them [23,24,40,41]. This observation suggests that other forces may counteract the membrane tension gradient for lipid flow. We hypothesize that the difference in membrane protein densities across the protein diffusion barriers or LTPs generates a new driving force for directional lipid flow termed lipid osmosis [11].

### Lipid osmosis and osmoc membrane tension

Osmosis plays a key role in the transport of water and other small molecules between different cell compartments, cells, tissues, and organs [42]. For example, our blood contains 60–80 mg/ml proteins, mainly human albumin proteins, that generate osmotic pressure to counteract hydrostatic blood pressure to draw extracellular fluid back into the blood and help to circulate ∼8 liters of fluid every day. Lipid osmosis is an analogous osmotic process occurring in two-dimensional membranes through nanoscale lipid filters. Biological membranes can be viewed as a two-dimensional fluid with lipids as a solvent and membrane proteins as solutes. When membrane proteins are enriched in a region with the help of a physical barrier or lipid filter that prevents the membrane proteins from diffusing into other membrane regions but allows the passage of lipids, lipids flow from the nearby regions to the protein-rich region due to lipid osmosis. To justify the process, we notice that the chemical potential of a lipid in a membrane subject to membrane tension *σ* is

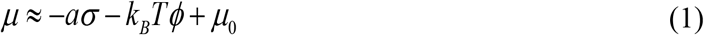

where *a* is the average lateral area per lipid, *k*_*B*_ is the Boltzmann constant, *T* is the absolute temperature with *k*_*B*_*T* =4.1 pN×nm at room temperature, *φ* ≪ 1 is the ratio of the number of mobile membrane proteins to that of lipids, and *μ*_0_ is the reference chemical potential independent of membrane tension and the protein-to-lipid ratio (see Eqs. S1-S5 in Supplementary Text for the derivation of Eq. 1) [43]. For simplicity, here we have assumed that there is only a single species of lipids in the membrane whose number dominates that of proteins. Moreover, the membrane proteins are mobile in the lipid bilayer and fully mixed with lipids. In other words, these proteins do not bind to the cytoskeleton, and neither proteins nor lipids form clusters. The chemical potential with multiple species of lipids and all ranges *φ* is shown in Eq. S10 in Supplementary Text. Thus, membrane tension and the presence of membrane proteins lower the chemical potential of lipids by decreasing their mechanical energy and increasing their mixing entropy, respectively.

To illustrate lipid osmosis, suppose an enclosed fence separates a planar membrane into two regions (Figure 2f). We assume that the membrane inside the fence has a mobile protein density higher than that of the outside, with a difference *c* ≈ Δ*φ*/*a*. The membrane tension inside and outside the fence is initially equal. At this moment, the lipids outside the fence have a higher chemical potential than that of the lipids inside. Consequently, lipids flow inside. As more lipids accumulate inside the fence, the inside membrane tension decreases, which gradually increases the lipid chemical potential and slows down the lipid flow. When the lipid transfer between the two membrane regions reaches thermodynamic equilibrium, the chemical potentials of lipids inside and outside the fence become equal, leading to lower membrane tension inside than outside, or the osmotic membrane tension

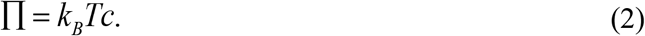

The density and composition of membrane proteins vary greatly among different biological membranes. For example, bacteria membranes and synaptic vesicles have a high protein density of ∼0.1 nm^-2^, while the plasma membranes of HeLa cells and kidney fibroblast contain 0.03–0.05 nm^-2^ integral proteins [31]. In contrast, autophagosomes bear few integral proteins [44]. Consequently, the osmotic membrane tension can be as high as 0.4 pN/nm, compared to the reported cell membrane tension of 0–0.4 pN/nm [23,32,39,45]. Therefore, a physiological protein density gradient can generate a high osmotic membrane tension. Correspondingly, each lipid in the membranes has a minimal chemical potential of -7% k_B_T assuming a typical lateral area of 0.7 nm^2^ for lipids. This magnitude of energy is equivalent to the potential energy of a water molecule at a water height of ∼1,000 meters! Therefore, lipid osmosis can modulate membrane tension and lipid flow, in addition to other well-known mechanical and chemical means, including forces produced by cytoskeleton and molecular motors, osmotic pressure, lipid synthesis or degradation, and membrane remodeling by proteins. Consequently, bulk lipid flow can be indirectly coupled to ATP hydrolysis or other energy sources via membrane tension or lipid osmosis. Finally, the osmotic membrane tension expressed by Eq. (2) assumes that lipid flow reaches thermodynamic equilibrium. However, given the highly dynamic nature of cell membranes, achieving true equilibrium may necessitate rapid lipid flow through lipid filters - a condition that might not always hold. Consequently, the instantaneous osmotic membrane tension in certain cell membrane regions could deviate from the predicted values.

Lipid osmosis and osmotic membrane tension may occur between different membranes mediated by bridge LTPs (Figure 1a), because the above thermodynamic analysis does not depend on the shape, size, and other details of lipid filters and membrane proteins. In general, the rate of the lipid flow *J* from a donor membrane to an acceptor membrane through a fence or a bridge LTP is proportional to the sum of the difference in the canonical membrane tension and the osmotic membrane tension, i.e.,

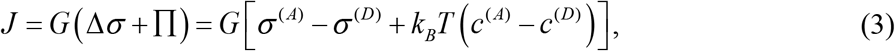

where the superscript indicates membranes (A = acceptor; D = donor) and *G* is the lipid permeability of the fence or bridge LTP [11]. This equation is a direct extension of the Starling equation governing the extracellular fluid flow between blood and tissues, where canonic and osmotic membrane tension replaces hydrostatic and colloid osmotic pressure between plasma inside microvessels and their outside interstitial fluid, respectively [46]. Equation (3) indicates that the gradients of membrane tension and the protein density may drive lipid flow synergistically or antagonistically, depending upon the signs of *σ*^(*A*)^ − *σ*^(*D*)^ and *c*^(*A*)^ − *c*^(*D*)^. When the two terms have opposite signs, little lipid flow or even reverse lipid flow may occur despite the presence of a significant membrane tension gradient. The latter scenario is analogous to reverse osmosis utilized in sea water desalting [42] and thus may be called reverse lipid osmosis.

### Membrane curvature, potential, and other factors in lipid flow

The chemical potential of lipids in a membrane also depends on the membrane curvature, the electric potential for charged lipids, the volume for an enclosed membrane, and lipid-lipid and lipid-protein interactions, thereby affecting lipid transfer. The bending energy of a membrane can be calculated by the Helfrich equation or other theories of membrane elasticity, which also depends on the global shape, spontaneous curvature, membrane tension, as well as the interior pressure of an enclosed membrane [47-49]. Here, we calculated the chemical potential of a lipid in a spherical vesicle due to membrane bending as

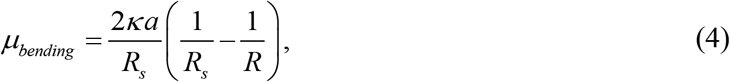

where *R* is the radius of the vesicle, *R*_*s*_ is the radius of spontaneous curvature of the membrane, and *κ* is the membrane bending rigidity with a typical value of ∼100 pN×nm (see Section 3 in Supplementary Text for the derivation of the equation). For small vesicles with a high spontaneous curvature, the chemical potential of its lipids can be high. For example, with *R*_*s*_ = 50 nm, the chemical potential is −9 ×10^−3^ k_B_T for *R* = 30 nm and 7 ×10^−3^ k_B_T for *R* = 100 nm. However, for vesicles whose membranes bear no spontaneous curvature (*R*_*s*_ → +∞), the chemical potential of their lipids due to membrane bending is zero regardless of the vesicle size [50].

The chemical potential of charged lipids strongly depends on the electric potential of the membrane or leaflet where the lipids are located, with *μ*_*electric*_ = *qV* where *q* is the charge per lipid and *V* is the electric potential of the membrane. LTP-mediated lipid transfer has been observed between ER and all other organelles with different lumen potential (Figure 1a). While the ER lumen is approximately neutral relative to the cytosol, the lumen of mitochondria or lysosome has an electric potential of −200 − −100 mV or +30 − +120 mV, respectively [51]. Even a small gradient of electric potential between different organelle membranes could provide a huge driving force for lipid transfer with the help of lipid scramblases. For example, the chemical potential of a single lipid with a unit charge in two membrane leaflets with a potential difference of 100 mV differs by 3.9 k_B_T, which is orders of magnitude greater than the chemical potential induced by physiological membrane tension and membrane curvature described above (10^−4^ −10^−2^ k_B_T).

It remains a puzzle how organelles or membranes maintain their stabilities by controlling the lipid flow under such high electric potential. Lipid transfer may be coupled to ion transduction across membranes, which may lower the chemical potential of lipids. Alternatively, lipid transfer may only occur when the membranes are transiently depolarized with minimal membrane potential. Supporting these views, many scramblases, including TMEM16F, are also ion channels and transfer both ions and lipids upon activation [17,52,53]. Finally, charged membrane proteins can help equilibrate charged lipids in the presence of a membrane potential gradient due to the Gibbs-Donna effect [46]. More work is required to distinguish the different scenarios of lipid transfer between polarized membranes.

The chemical potential of a lipid strongly depends on its membrane environment due to lipid-lipid and lipid-protein interactions. For example, cholesterol tends to flow to membranes containing lipids with long saturated acyl chains [54]. Finally, the lipid transfer associated with the membrane expansion of an organelle is often accompanied by its volume change, which may change the chemical potential of lipids in the membrane, thereby facilitating or opposing lipid transfer depending on the pressure difference on the two sides of the membrane. In conclusion, the chemical potential of a lipid critically depends on its environment, including membrane tension, protein density, electric potential, volume of an enclosed membrane, and lipid-lipid and lipid-protein interactions. These factors together determine the direction and amount of lipid flow. In the following, we use the lipid chemical potential to quantify the lipid transfer mediated by LTPs in the default presence of scramblases.

### Bridge LTPs

Suppose a bridge LTP transfers lipids between a donor membrane and an acceptor membrane in the presence of gradients of both membrane tension and potential, with other factors affecting lipid flow being approximately equal for both membranes. Then upon equilibrium, the fractions of any species of lipids 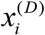 and 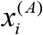 in the donor and acceptor membranes, respectively, satisfy the following Boltzmann distribution derived from equal chemical potentials of lipids in the two membranes (see Section 2.1 of Supplementary Text for its derivation)

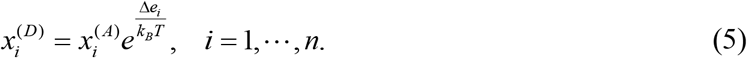

Here, the subscript *i* designates the *i*-th lipid species in a total of *n* different lipid species present in both membranes, and

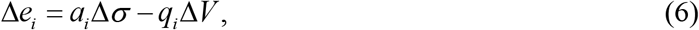

Where Δ*σ* = *σ*^(*D*)^ − *σ*^(*A*)^ and Δ*V* = *V*^(*D*)^ − *V*^(*A*)^ are the differences in membrane tension and potential, respectively, and *a*_*i*_ and *q*_*i*_ are the lateral area and charge of the lipid, respectively. Therefore, the relative lipid distributions strongly depend on the membrane tension difference, as well as the potential difference for charged lipids. Consequently, lipids flow to the membrane with a high membrane tension and a low membrane potential for charged lipids (Figures 1a, 3a, and 3c). The absolute distribution of each species of lipids in each membrane can be derived from Eq. (5), as is shown by Eqs. S24-S26 in Supplementary Text. These derivations highlight the role of lipid osmosis in membrane expansion, which can be better seen in the absence of any membrane tension and potential gradients, or Δ*e*_*i*_ = 0, *i* = 1,⋯, *n*. In this case, lipids are drawn to the membrane with a high protein density (Eq. S27). Upon equilibrium, the fractions of proteins in different membranes become identical, and so do the fractions of each lipid species (Figure 3b and Eq. S28). Therefore, the equilibrium size of the membrane is approximately proportional to the number of membrane proteins in the membrane, and all lipids are fully mixed. However, even though the equilibrium size of the membrane is altered by the membrane tension gradient, all lipids remain fully mixed (Figure 3c and Eq. S31).

**Figure 3.**
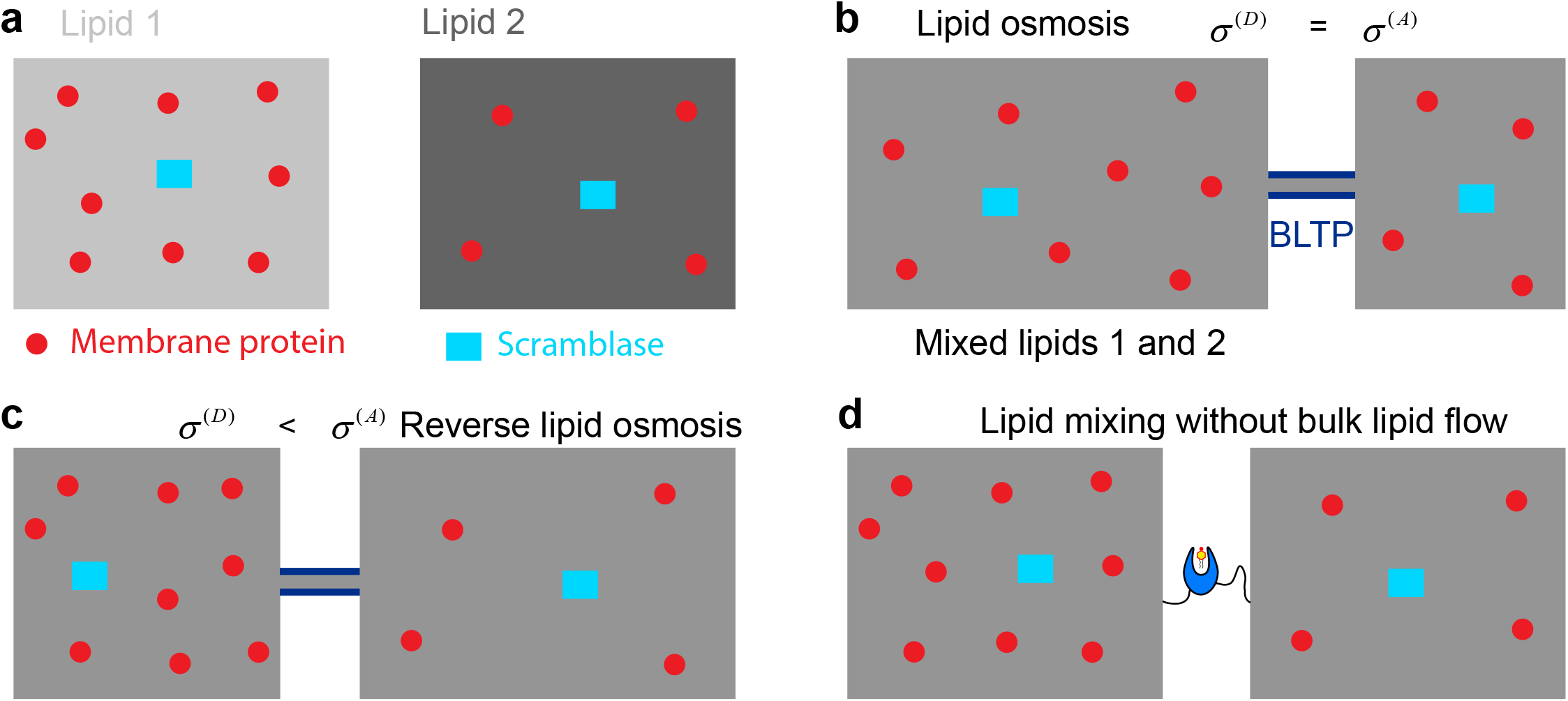
Areas and lipid distributions of two membranes before and after the LTP-mediated lipid transfer reaches equilibrium. **(a)** The two membranes initially have identical areas but contain different lipid species (indicated by different grey scales) and densities of membrane proteins (red circles and cyan rectangles) before lipid transfer. **(b)** In the absence of membrane tension gradient, lipid osmosis through BLTPs causes expansion of the membrane with a higher density of membrane proteins, leading to equal densities of membrane proteins and fractions of lipids in both membranes upon equilibrium. **(c)** A high gradient of membrane tension counteracts lipid osmosis to expand the membrane with lower membrane protein density, leading to reverse lipid osmosis. **(d)** Lipid transfer mediated by shuttle LTPs causes lipid mixing without changes in membrane areas in a manner independent of the protein density gradient.

The rate of bulk lipid flow through a bridge LTP appears to be high in vivo (>100 s^-1^) [55] but has not been accurately measured [11]. To estimate the flow rate, we approximate the LTP as a lipid bridge with a width *d* and a length *l* between two membranes with a membrane tension difference Δ*σ* and potential difference Δ*V* (Figure 1a). Treating the lipid bilayer as a two-dimensional continuous and viscous fluid, we solved the Navier-Stokes equation governing the fluid dynamics, yielding the unidirectional bulk lipid flow rate through a single bridge LTP as

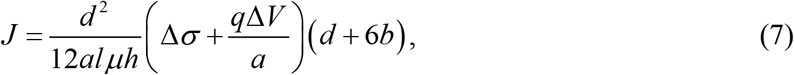

where *μ* and *h* are the viscosity and thickness, respectively, of the lipid bilayer, *q* is the average charge per lipid, and *b* is the slip length characterizing the adhesion of lipids to the BLTP groove (see Section 5 of Supplementary Text for its derivation). Equation (7) is essentially the two-dimensional version of the Hagen-Poiseuille equation [43]. The rate strongly depends on the groove width of bridge LTPs, which varies from 1 nm to 3 nm along their lengths based on the AlphaFold structures of a few bridge LTPs without lipids in their grooves [56] (Figure 1c). For simplicity, we chose an average width of *d* = 2 nm for our estimation. In addition, we adopted a sticky boundary condition with *b* = 0. Assuming *μh* ≈ 5.8×10^−7^ pN× s/nm, *a* = 0.7 nm^2^, *l* = 20 nm, Δ*V* = 0, and Δ*σ* = 0.001–0.1 pN/nm, we estimate the lipid flux *J* = 80 − 8208 s^-1^. Similarly, in the presence of 50 mV potential gradient and an average of 0.1 unit charge per lipid, but no membrane tension gradient, the lipid flux is ∼10^5^ s^-1^. In the presence of lipid osmosis or a protein density difference between the two membranes, the membrane tension difference Δ*σ* in Eq. (7) should be replaced by Δ*σ* +∏, as is indicated by Eq. (3). Thus, physiological membrane tension or potential gradient can drive rapid bulk lipid flow consistent with in vivo observations [55].

The limitations of our above estimation must be acknowledged. While the Navier-Stokes equation has found widespread applications in characterizing fluid flow at the nanoscale [57], particularly the water transport through aquaporin [58], its applicability to lipid flow through the BLTP remains to be validated, and the slip length needs to be measured. Encouragingly, the calculated lipid flux is close to our previous estimation based on single-file lipid transport through the BLTP [11], providing further evidence of rapid lipid flow. Both estimations assumed that lipids can move freely from the donor membrane into the BLTP and out of the BLTP to the acceptor membrane. However, lipids may encounter significant energy barriers during such movement, potentially limiting the overall lipid flow rate. To address these uncertainties, a reconstituted assay for bulk lipid transfer in vitro would be invaluable (more discussion in the last section).

The unidirectional bulk lipid flow differs from simple lipid diffusion through LTPs, which are bidirectional without bulk lipid flow (Figure 3d). In a typical experiment to test lipid transfer, a lipid transfer rate is often derived from the measured rate of fluorescent tracer lipids diffusing from one membrane to the other through LTPs. Given the same model for the bridge LTP (Figure 1a), the rate can be derived as

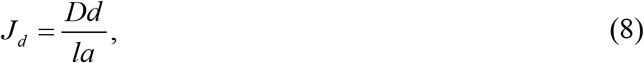

where *D* is the lateral diffusion constant of the lipid (see Section 6 of Supplementary Text). Choosing *D* = 5 μm^2^/s and other parameters as above, we estimated the lipid transfer rate as *J*_*d*_ = 3.6×10^5^ s^-1^. Thus, lipids can rapidly diffuse through a bridge LTP without bulk lipid flow.

### Shutle LTPs

Suppose a shuttle LTP transfers two species of lipids designated as 1 and 2 in the subscript between a donor membrane and an acceptor membrane through a lipid exchange mechanism (Figure 1c). While bridge LTPs transport many if not all species of lipids, shuttle LTPs only transfer their cognate lipids [4,10,54]. When the lipid transfer reaches equilibrium, the lipid fractions of the two lipid species in the two membranes satisfy the following equation (see Section 2.2 of Supplementary Text for its derivation)

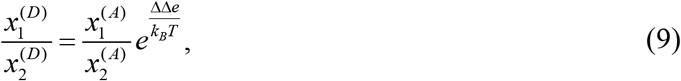

where

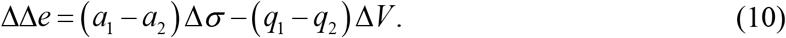

The absolute distributions of lipids in both membranes can also be derived based on Eq. (9). Both equations suggest that the equilibrium lipid distributions are sensitive to the membrane tension gradient only when the two lipid species differ in their lateral areas (or *a*_1_ ≠ *a*_2_). For example, Osh4p exchanges cholesterol and PI4P with a significant difference in their lateral areas (0.45 nm^2^ for cholesterol and ∼0.7 nm^2^ for PI4P) [9,54]. Therefore, Osh4p-mediated lipid transfer is predicted to be sensitive to membrane tension and more PI4P is drawn to membranes with higher membrane tension. However, shuttle LTPs are generally less sensitive to membrane tension gradient in lipid transfer than bridge LTPs are. In contrast, membrane potential gradient is expected to greatly affect lipid transfer by shuttle LTPs, such as Osh4p, because their cognate lipids often differ in charge. Moreover, shuttle LTPs do not cause bulk lipid transfer between membranes because the lipid exchange scheme only alters the lipid composition but not the total number of lipids in each membrane (Figure 3d). Consequently, the equilibrium lipid distributions are independent of protein distributions in both membranes. In other words, lipid osmosis does not play a role in the equilibrium lipid distributions.

The rate of lipid transfer mediated by shuttle LTPs has been solved analytically or numerically [11]. The rate depends on two lipid exchange rate constants associated with each membrane (Figure 1c). The rate constants for different membranes are generally different, corresponding to ΔΔ*e* ≠ 0. If the two rate constants are the same for both donor and acceptor membranes, or 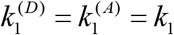 and 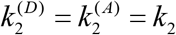, corresponding to the case of ΔΔ*e* = 0, and the LTP is free in solution without a membrane tether, the rate to transfer both species of lipids can be expressed as

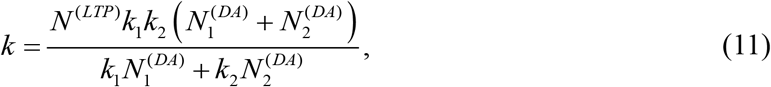

Where *N* ^(*LTP*)^ is the total number of shuttle LTPs and *k*_1_ and *k*_2_ are the exchange rate constants leading to the extraction of lipid species 1 and 2, respectively, by the LTP. This formula has been used to fit the measured rates of the lipid transfer mediated by Osh4p [11].

### Lipid osmosis between two leaflets

Lipid osmosis is also expected to occur between two leaflets of a membrane through scramblases. Lipid scramblases contain a hydrophilic groove or thin the surrounding membrane to facilitate lipids to flip between the two leaflets [16,52,53]. The two leaflets often bear different densities of peripheral membrane proteins, causing lipid osmosis to draw more lipids to the leaflet with a higher density of proteins. The accumulation of lipids generates membrane curvature in a manner dependent on the protein density gradient. To appreciate the role of lipid osmosis in membrane bending, we modified the area-difference-elastic model of membranes [49] to include the effect of lipid osmosis and derive the curvature induced by lipid osmosis (see Section 4 of Supplementary Text) as

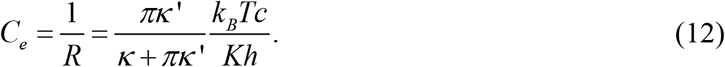

Here, *R* is the radius of curvature of the membrane, *κ* and *κ* ‘are the local and non-local membrane bending rigidities, respectively, *K* is stretching modulus of the bilayer, and *c* is the difference in the densities of peripheral proteins in the two leaflets. This equation indicates that the equilibrium membrane curvature due to lipid osmosis is proportional to the difference in the protein densities in both leaflets. The stretch modulus is 100 − 500 pN/nm for purified membranes and as small as 0.04 pN/nm for biological membranes, while for typical membranes, *κ* ≈ *κ*’, *h* = 2.5 nm [24,31,49,59]. With these membrane parameters, we estimated a protein density gradient of *ρ* =0.1 nm^-2^ bends the membrane with a radius of curvature of 4 − 2000 nm with a stretch modulus ranging from 0.5 pN/nm to 250 pN/nm. Thus, lipid osmosis across two leaflets of membranes can generate significant membrane curvature by driving scramblase-mediated lipid flow.

### Potential roles of lipid osmosis in membrane dynamics

Lipid osmosis tends to draw lipids to membrane domains with higher protein densities, thereby expanding the membranes. Therefore, lipid osmosis likely plays an important role in the biogenesis and maintenance of many organelles associated with membrane expansion or replenishment. BLTPs are involved in the membrane expansion of phagophores, lipid droplets, prospores during yeast sporulation, acrosome in sperms, and Gram-negative bacteria [8,60]. Various proteins coat the necks between different membrane domains (Figure 2). These BLTPs and protein coats prevent membrane proteins from diffusing away from new organelles, which may facilitate membrane expansion or lipid homeostasis. Growing organelles, such as phagophores and lipid droplets, are often crowded with membrane proteins on their surfaces, especially peripheral proteins recruited from the cytosol [37,38] (Figures 2b & 2c). For example, an average of 272 Atg8 proteins are covalently attached to PE lipids on a phagophore, which is crucial for its growth [61] (Figure 1d). On a phagophore with a radius of 30 nm, the Atg8 protein alone contributes a protein density of 0.024 nm^-2^, corresponding to a high osmotic membrane tension of 0.1 pN/nm. Note that many of these proteins play additional roles in organelle biogenesis, such as membrane remodeling and recruitment of other proteins. Finally, overexpression of many membrane proteins generates inducible intracellular membranes, which appear to be artificial membranes with sizes ranging from tens of nanometers to several micrometers and different shapes [62,63]. They can bud from different organelles where the membrane proteins are over-expressed and are observed in bacteria, yeast, and mammalian cells. Often, the transmembrane domains of these proteins alone are sufficient for forming new membrane buds, regardless of their cytosolic domains or enzymatic activities. All these observations are consistent with the potential role of lipid osmosis in membrane expansion.

Lipid osmosis can be strong enough to overcome other forces opposing organelle growth, such as membrane bending. Based on Eq. (1), the total lipid-protein mixing energy of a membrane can be estimated as −*N*_*p*_*k*_*B*_*T*. Therefore, the presence of each mobile membrane protein contributes to an energy of *k*_*B*_*T* favoring membrane growth. Assuming a spheric vesicle growing from a relaxed protein-free membrane (Figure 2b), its total bending energy is 8*πκ* or ∼500 *k*_*B*_*T* regardless of its radius. Therefore, ∼500 membrane proteins are sufficient to overcome the bending energy to stabilize the growing organelle. Assuming a maximum protein density of 0.2 nm^-2^ due to size exclusion, we estimated a minimum radius of 14 nm for the organelle. The estimation suggests that lipid osmosis alone is strong enough to counteract membrane bending to support a growing vesicle with a radius greater than 14 nm. The initial growth below such a small radius probably needs membrane bending activities of proteins and/or mechanical forces provided by cytoskeleton or molecular motors.

Osmotic membrane tension may play an important role in organelle biogenesis. Nascent organelles generally have a smaller radius of membrane curvature (*R*) than their connected membranes, while both membranes experience the same interior pressure due to their connected lumens (Figure 2). For the two membranes to co-exist with the same pressure (Δ*p*), the more curved membrane requires lower membrane tension (*σ*), as suggested by the Young-Laplace equation (*σ* = Δ*p* × *R*/2) [64]. Osmotic membrane tension can stabilize the nascent organelles by lowering their membrane tension in a manner dependent upon the membrane protein density, which may facilitate the formation of nascent vesicles or lipid droplets (Figures 2b & 2c). A more accurate and quantitative assessment of lipid osmosis shall consider the shape, membrane tension, pressure, and other factors of the organelle [47,48,65].

### Methods to test lipid osmosis and lipid transfer

Despite the potential of bridge LTPs to mediate fast bulk lipid flow observed in the cell [66], a few bridge LTPs tested so far can only slowly transfer lipids between vesicles in vitro with a few lipids per minute per LTP [2,67,68]. The lipid transfer probably occurs via a shuttle mechanism in the absence of net driving forces for lipid flow. Therefore, it is imperative to reconstitute the fast bulk lipid flow through bridge LTPs and test the various driving forces, especially lipid osmosis, described in the preceding sections. However, these driving forces often counteract each other to oppose fast lipid flow. For example, bulk lipid flow would cause a change in the membrane area of a vesicle, which in turn causes changes in membrane tension and curvature if the membrane tension is not controlled or the volume of the vesicle is kept constant. As a result, bulk lipid flow is unlikely to occur between small unilamellar liposomes (SUVs), as previously observed [67]. Therefore, it is crucial to study lipid flow in a system where each of the parameters affecting lipid flow can be independently controlled. In contrast to cell membranes, model membranes allow fine-tuning of the lipid composition, curvature, tension, and separation of membranes. Additionally, the model membranes generally allow the incorporation of proteins that regulate membrane flow, including those acting as fences, bridges, and shuttles. For proteins that are difficult to purify or reconstitute, their functions may be surrogated by rationally designed nanostructures. In this section, we review a few tools that may be used to characterize the energetics and kinetics of lipid flow in a cell-free environment, with an emphasis on the utilities of cutting-edge membrane engineering methods enabled by DNA nanotechnology [69].

Several commonly used model membranes are suitable for experimentally studying lipid flow, including unilamellar liposomes (SUVs and GUVs), supported bilayers, and free-standing bilayers [70-73]. For example, to examine lipid transfer across two apposed membranes, one can potentially bring two liposomes (or a liposome and a supported bilayer) containing reconstituted LTPs into proximity. Fluorescent tracers embedded in membranes can be used to track lipid flow, and the bulk lipid flow resulting in the growth of GUVs may be directly observed under microscopy. *<u>Tension</u>* of the two membranes can be tuned by osmolarity pressure and/or micropipette aspiration (**Figure 4a**) while being monitored by tension-sensitive fluorophores such as FliptR [74] or membrane tethers pulling [75]. Chemically modified lipids can be incorporated into the membranes for subsequent recruitment of proteins to change the *<u>protein density</u>*. Membrane *<u>curvatures</u>* can be controlled by using liposomes of well-defined radii (from tens to several hundred nanometers; see our theoretical discussion above). Conventional methods (e.g., extrusion) often lead to heterogeneous liposome sizes that are difficult to control deterministically. In contrast, DNA cages carrying amphipathic moieties have served as molds for membrane reconstitution to generate sub-100 nm liposomes with predefined, uniform sizes (**Figure 4b**, left) [76]. Attaching DNA bricks to heterogeneous liposomes has enabled the centrifugation-based size sorting (30–150 nm) that led to liposome populations with very narrow size distributions [77]. The *<u>inter-membrane distance</u>* can be controlled with nanometer precision by rigid DNA segments anchored on both membranes [78-80] (**Figure 4b**, right), mimicking membrane contact sites in cells. For example, liposome pairs held at controlled distances have helped shed light on the mechanism of lipid-transferring protein E-Syt1, suggesting its shuttle-like transport activities are dependent on its membrane tether length relative to the inter-membrane distance [79].

**Figure 4.**
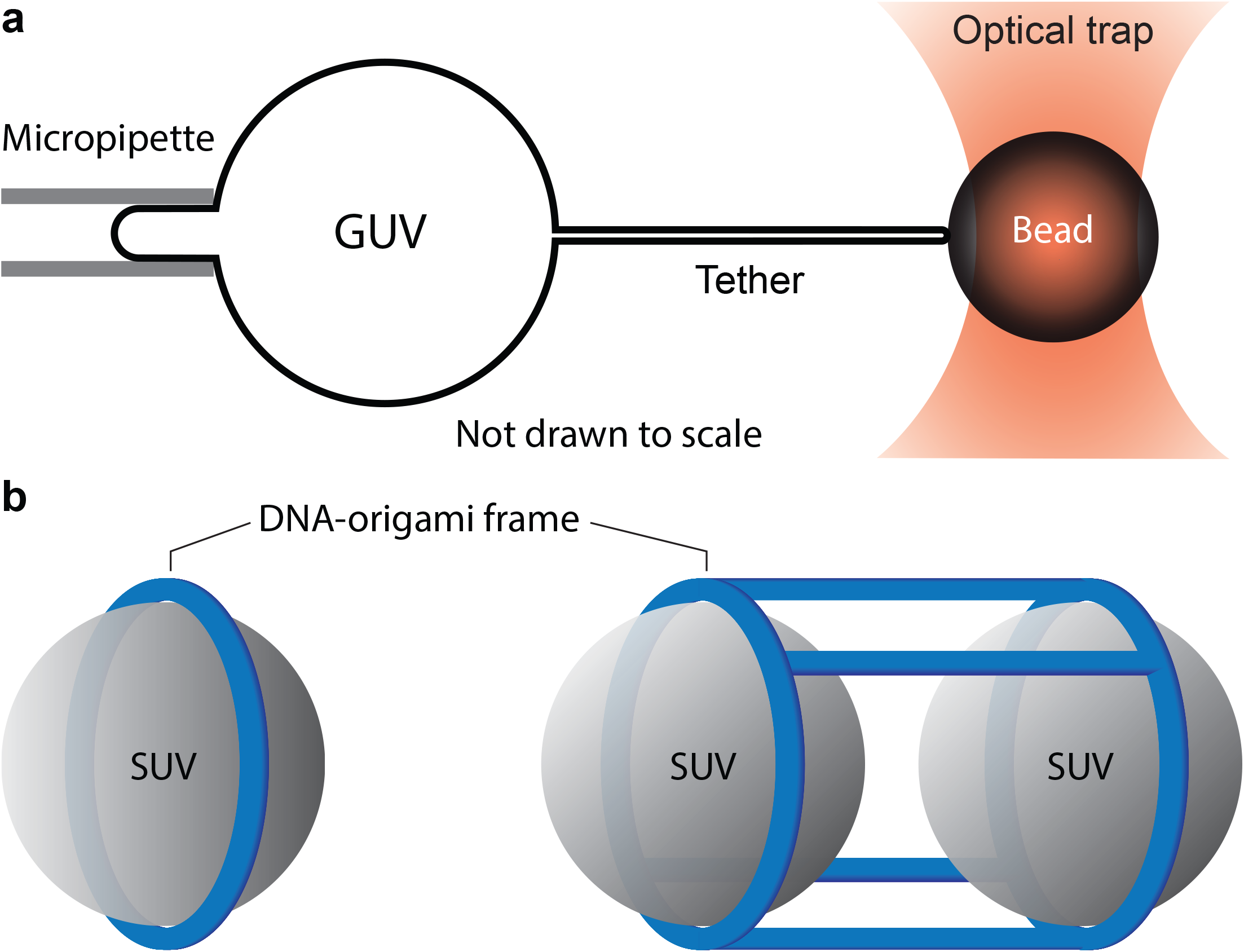
Methods for controlling tension, curvature, and separation of model membranes. **(a)** Membrane tension of a GUV can be controlled by micropipette aspiration and measured by membrane tether pulling with optical tweezers. Different suction pressures can be applied via the micropipette to change membrane tension. The pulling force measured by the optical trap can be used to calculate the membrane tension. **(b)** Membrane curvature and inter-membrane distance of SUVs (grey) can be controlled by DNA-origami frames (blue). These DNA nanostructures can be modified with amphipathic molecules on the interior (not shown for clarity) and serve as templates for vesicle reconstitution. Left: a size-defined SUV formed inside a DNA ring. Right: a pair of SUVs separated by a set of rigid DNA pillars with defined lengths.

To study the lateral lipid flow within the same membrane, it is conceivable to use a GUV with its membrane laced with cytoskeleton and membrane protein fences. Although it is possible to reconstitute dynamic actin networks on model membranes [81,82], the membrane domains defined by the actin fences are usually irregular and hard to finetune, making it difficult to manipulate and measure the properties of each domain. Therefore, one may consider using membrane-anchored DNA filaments as an artificial cytoskeleton [83]. Indeed, various curved or straight DNA filaments have been assembled on or inside GUVs [84-88]. By programming the assembling pattern of the DNA filaments and their membrane anchors, one could build a series of picket-and-fence like architectures that limit in-plane protein (but not lipid) diffusion, making it possible to generate a protein-density gradient and test lipid osmosis.

The state-of-the-art membrane manipulation methods offer a promising solution to reconstituting lipid flow in a cell-free environment. However, several technical challenges remain. For example, independently tuning the tension of two or multiple membranes in a system is likely necessary to generate lipid flow, yet difficult to achieve. More versatile and user-friendly tension-sensing probes are also needed to monitor the mechanical stress of membranes in real-time. These challenges call for novel molecular tools that directly address the mechanical properties of membranes and manage leaflet homeostasis. We propose that because DNA nanostructures can undergo conformational changes to exert force to drive membrane deformation on demand [89], it should be possible to build DNA-based devices that modulate membrane tension in a stepwise and targeted manner. Similarly, mechanosensitive DNA structures embedded in membranes, such as a transmembrane channel with dilatable diameter, can serve as membrane tension sensors [90]. Coincidentally, DNA nanopores can facilitate molecular exchange across membranes and scramble lipids between leaflets [91].

## Acknowledgments

We thank Avinash Kumar for reading the manuscript and Jie Yang for figure preparation. YZ derived the formulae and YZ and CL wrote the manuscript. This work was supported by National Institutes of Health grants R35 GM131714 (YZ) and R35GM149264 (CL).

