## Supplementary Text for "Lipid osmosis, membrane tension, and other mechanochemical driving forces of lipid flow"

### 1. Chemical potential of lipids due to membrane tension and lipid-protein mixing

Suppose a membrane of area $S$ contains $N$ identical lipids and $N_p$ membrane proteins and is subject to a membrane tension $\sigma$, then the free energy of the membrane based on equilibrium statistical mechanics can be written as

$$F = -\sigma S + k_B T\left[N \ln \frac{N}{N+N_p} + N_p \ln \frac{N_p}{N+N_p}\right] + N\mu_0 \tag{S1}$$

where the second term with a parathesis is the energy due to the entropy of mixing proteins and lipids, and $N\mu_0$ accounts for other energy contributions [1]. Note that the membrane area is

$$S = Na + N_p a_p, \tag{S2}$$

where $a$ and $a_p$ are the average lateral area per lipid and protein, respectively. The chemical potential of the lipid is

$$\mu = \frac{\partial F}{\partial N} = -a\sigma + k_B T \ln \frac{N}{N+N_p} + \mu_0. \tag{S3}$$

For simplicity, suppose the number of lipids is much greater than that of the membrane protein, then

$$\phi \equiv \frac{N_p}{N} \ll 1. \tag{S4}$$

Under this condition, Eq. (S3) can be simplified as

$$\mu \approx -a\sigma - k_B T\phi + \mu_0. \tag{S5}$$

This equation also gives the chemical potential energy per membrane area as

$$\begin{aligned} \mu_a &\approx \frac{\mu}{a} = -\sigma - k_B T\frac{\phi}{a} + \frac{\mu_0}{a} = -\sigma - k_B T\frac{N_p}{Na} + \frac{\mu_0}{a} \\ &\approx -\sigma - k_B Tc + \frac{\mu_0}{a} \end{aligned} \tag{S6}$$

where $c$ is the number of membrane proteins per unit membrane area. The chemical potential of lipids in the absence of membrane proteins was previously derived based on an ensemble of constant area per lipids [2], while our derivation assumes an ensemble of constant membrane tension. Results derived from both ensembles are consistent, given the high stretching modulus of membranes.

If a membrane contains many species of lipids in addition to membrane proteins, then the free energy shown in Eq. (S1) becomes

$$F = -\sigma S + k_B T\left(\sum_{i=1}^{n} N_i \ln \frac{N_i}{N_t} + N_p \ln \frac{N_p}{N_t}\right), \tag{S7}$$

where $N_i, i=1,\cdots,n$ is the number of lipids of the *i*-th species, and $N_t$ is the total number of lipids and membrane proteins and other energy contributions are neglected, or

$$N_t = N_p + \sum_{i=1}^{n} N_i, \tag{S8}$$

and

$$S = N_p a_p + \sum_{i=1}^{n} N_i a_i, \tag{S9}$$

where $a_i$ is the average lateral area of the lipid species $i$. The chemical potential of the lipids of the *i*-th species is calculated as

$$\mu_i = \frac{\partial F}{\partial N_i} = -a_i\sigma + k_B T \ln \frac{N_i}{N_t}. \tag{S10}$$

### 2. Energetics of lipid transfer or through LTPs or barriers for protein diffusion

In the presence of a membrane potential, the chemical potential of a lipid species $i$ in a membrane containing a mixture of different species of lipids and proteins can be written as

$$\mu_i = -a_i\sigma + q_i V + k_B T \ln x_i, \tag{S11}$$

where $q_i$ is the charge of the lipid species $i$, $V$ is the membrane potential, and $x_i$ is the number fraction of the lipid species $i$, i.e.,

$$x_i \equiv \frac{N_i}{N_t}. \tag{S12}$$

Other factors that affect the chemical potential of lipids are assumed to be independent of lipid species and membranes. Suppose two membranes, a donor membrane and an acceptor membrane, designated as D and A in parenthesis in the superscript, respectively, contain multiple species of lipids as well as membrane proteins. The number of lipid species $i, i=1,\cdots,n$ and the total number of proteins in a membrane $M$ are designated as $N_i^{(M)}$ and $N_p^{(M)}$, respectively, with $M = D$ for the donor membrane and $M = A$ for the acceptor membrane. Note that lipids do not necessarily move from the donor membrane to the acceptor membrane, depending upon experimental conditions.

#### 2.1. Bridge LTPs and barriers that pass all lipid species

When lipid flow reaches equilibrium, the chemical potentials of each species of lipids in the two membranes become equal, i.e.,

$$\mu_i^{(D)} = \mu_i^{(A)}, \quad i=1,\cdots,n \tag{S13}$$

Substitution of Eq. (S11) into Eq. (S13), we have

$$a_i\Delta\sigma - q_i\Delta V = k_B T \ln \frac{x_i^{(D)}}{x_i^{(A)}}, \quad i=1,\cdots,n. \tag{S14}$$

Eq. (S14) can be rewritten as

$$x_i^{(D)} = x_i^{(A)} e^{\frac{\Delta e_i}{k_B T}} = x_i^{(A)} \big/ \beta_i, \quad i = 1, \cdots, n, \tag{S15}$$

where

$$\begin{aligned} \Delta e_i &= a_i \Delta\sigma - q_i \Delta V, \\ \beta_i &= e^{-\frac{\Delta e_i}{k_B T}}, \end{aligned} \tag{S16}$$

with

$$\begin{aligned} \Delta\sigma &= \sigma^{(D)} - \sigma^{(A)}, \\ \Delta V &= V^{(D)} - V^{(A)}. \end{aligned} \tag{S17}$$

Equation (S15) describes the requirement for the lipid transfer to reach equilibrium under all conditions. When the membrane tension and potential remain constant during lipid transfer, the lipid distribution of each lipid species is adjusted to satisfy Eq. (S15). In contrast, if the membrane tension or potential can vary during the lipid transfer process, the lipid distribution varies with membrane tension or potential as predicted by the equation.

The equilibrium fractions of all lipid species in each membrane can be solved from Eq. (S15) given experimental conditions or control parameters. The total number of lipids of the *i*-th species in both membranes $N_i^{(DA)}$ is a constant during lipid transfer, which yields

$$N_t^{(D)} x_i^{(D)} + N_t^{(A)} x_i^{(A)} = N_i^{(DA)}, \quad i = 1, \cdots, n, \tag{S18}$$

where $N_t^{(D)}$ and $N_t^{(A)}$ are the total numbers of all lipids and membrane proteins in the donor membrane and the acceptor membrane, respectively, *under the equilibrium condition*. Substituting Eq. (S15) into Eq. (S18), we have

$$\begin{cases} x_i^{(D)} = \dfrac{N_i^{(DA)}}{N_t^{(A)} \beta_i + N_t^{(D)}}; \\ x_i^{(A)} = \dfrac{N_i^{(DA)} \beta_i}{N_t^{(A)} \beta_i + N_t^{(D)}}. \end{cases} \tag{S19}$$

Suppose the numbers of membrane proteins in the donor membrane and the acceptor membrane are $N_p^{(D)}$ and $N_p^{(A)}$, respectively, we have

$$\begin{cases} N_t^{(D)} \sum_{i=1}^{n} x_i^{(D)} + N_p^{(D)} = N_t^{(D)}, \\ N_t^{(A)} \sum_{i=1}^{n} x_i^{(A)} + N_p^{(A)} = N_t^{(A)}. \end{cases} \tag{S20}$$

Substituting Eq. (S19) into Eq. (S20), we have

$$\begin{cases} N_t^{(D)} \left( 1 - \sum_{i=1}^{n} \dfrac{N_i^{(DA)}}{N_t^{(A)} \beta_i + N_t^{(D)}} \right) = N_p^{(D)}, \\ N_t^{(A)} \left( 1 - \sum_{i=1}^{n} \dfrac{N_i^{(DA)} \beta_i}{N_t^{(A)} \beta_i + N_t^{(D)}} \right) = N_p^{(A)}. \end{cases} \tag{S21}$$

The numbers of membrane proteins in the donor and acceptor membranes, or $N_p^{(D)}$ and $N_p^{(A)}$ remain constant during lipid transfer and are given as experimental control parameters. If the differences in membrane tension and potential do not change during lipid transfer, the energy differences $\Delta e_i, i=1,\cdots,n$ are constants. Therefore, the total numbers of lipids and membrane proteins in equilibrium, or $N_t^{(D)}$ and $N_t^{(A)}$ can be solved from Eq. (S21). To facilitate their computations, we scaled all the lipid and protein numbers by their total number in both membranes, or

$$N_t^{(DA)} = \sum_{i=1}^{n} N_i^{(DA)} + N_p^{(D)} + N_p^{(A)} = N_t^{(D)} + N_t^{(A)}, \tag{S22}$$

yielding

$$\begin{aligned} &y_t^{(D)} = \frac{N_t^{(D)}}{N_t^{(DA)}} = 1 - y_t^{(A)}; \quad y_t^{(A)} = \frac{N_t^{(A)}}{N_t^{(DA)}} = 1 - y_t^{(D)}; \\ &y_p^{(D)} = \frac{N_p^{(D)}}{N_t^{(DA)}}; \quad y_p^{(A)} = \frac{N_p^{(A)}}{N_t^{(DA)}}; \quad y_p^{(DA)} = y_p^{(D)} + y_p^{(A)}; \quad y_i^{(DA)} = \frac{N_i^{(DA)}}{N_t^{(DA)}}. \end{aligned} \tag{S23}$$

Then Eq. (S21) can be simplified as

$$y_t^{(D)} \left( 1 - \sum_{i=1}^{n} \frac{y_i^{(DA)}}{(1-\beta_i) y_t^{(D)} + \beta_i} \right) = y_p^{(D)}. \tag{S24}$$

We can numerically solve $y_t^{(D)}$ and calculate other distributions as

$$\begin{cases} x_i^{(D)} = \dfrac{y_i^{(DA)}}{(1-\beta_i) y_t^{(D)} + \beta_i}; \\ x_i^{(A)} = \dfrac{y_i^{(DA)} \beta_i}{(1-\beta_i) y_t^{(D)} + \beta_i}. \end{cases} \tag{S25}$$

In addition, we can calculate the distribution of each species of lipids in either of the membrane as

$$z_i^{(M)} = \frac{N_i^{(M)}}{N_i^{(DA)}} = \frac{x_i^{(M)} y_t^{(M)}}{y_i^{(DA)}}; \quad M = D, A; \quad i = 1,\cdots,n. \tag{S26}$$

To better appreciate the role of lipid osmosis in the lipid flow, we consider a simple case where the differences in membrane tension and potential between the two membrane vanish, or $\Delta e_i = 0$ and $\beta_i = 1, i=1,\cdots,n.$ In this case, we can solve for $y_t^{(D)}$ from Eq. (S24), yielding

$$y_t^{(M)} = \frac{y_p^{(M)}}{y_p^{(D)} + y_p^{(A)}} = \frac{N_p^{(M)}}{N_p^{(D)} + N_p^{(A)}}, \quad M = D, A. \tag{S27}$$

Similarly, we have

$$x_i^{(M)} = y_i^{(DA)}; \quad x_p^{(M)} = y_p^{(DA)}; \quad z_i^{(M)} = y_t^{(M)}; \quad M = D, A; \quad i = 1,\cdots,n. \tag{S28}$$

Equations (S27) and (S28) indicate that lipid osmosis causes a redistribution of lipids between different membranes to even the concentrations or fractions of all species of lipids and membrane proteins. As a result, the amount of lipids in each membrane is proportional to the protein amount in the membrane.

In another case where all lipids have the same lateral area and charge, but the two membranes have different membrane tension, corresponding to the case illustrated in Figure 3c, the parameter $\beta_i = \beta, i=1,\cdots,n$ is independent of lipid species. From Eq. (S25), we have

$$\begin{cases} \sum_{i=1}^{n} x_i^{(D)} = \dfrac{1-y_p^{(DA)}}{(1-\beta)\, y_t^{(D)}+\beta}; \\ \sum_{i=1}^{n} x_i^{(A)} = \dfrac{\left(1-y_p^{(DA)}\right)\beta}{(1-\beta)\, y_t^{(D)}+\beta}. \end{cases} \tag{S29}$$

Let's define the fraction of each lipid species among the total lipids in a membrane $M$ as

$$\lambda_i^{(M)} = \frac{N_i^{(M)}}{\sum_{i=1}^{n} N_i^{(M)}}, \quad M = D, A, \tag{S30}$$

then from Eq. (S29), we have

$$\lambda_i^{(M)} = \frac{y_i^{(DA)}}{1-y_p^{(DA)}}, \quad M = D, A. \tag{S31}$$

This equation indicates that upon equilibrium of lipid transfer, each lipid species is fully mixed among both membranes, leading to their equal fractions among membranes, despite that more lipids populate in the membrane with higher membrane tension (indicated by the same grey scale of both membranes but a bigger area of the right membrane shown in Figure 3c).

In conclusion, the equilibrium lipid distributions of lipids in both membranes can be determined by the equal chemical potential of their lipids and lipid osmosis and membrane tension play important roles in the equilibrium distributions.

### 2.2. Shuttle LTPs

Suppose a shuttle LTP transfers two species of lipids designated as 1 and 2 in the subscript between the two membranes (Figure 1c). For simplicity, we assume that the LTP is not tethered to membranes, as many LTPs do [3-6]. When the lipid transfer reaches equilibrium, the two lipid exchange reactions (Figure 1c) lead to the following two equations

$$\begin{cases} \mu_1^{(D)} + \mu_2^{(LTP)} = \mu_2^{(D)} + \mu_1^{(LTP)}, \\ \mu_1^{(A)} + \mu_2^{(LTP)} = \mu_2^{(A)} + \mu_1^{(LTP)}, \end{cases} \tag{S32}$$

where $\mu_1^{(LTP)}$ and $\mu_2^{(LTP)}$ are the chemical potentials of lipid 1 and lipid 2, respectively, bound by the LTP. Removing both chemical potentials from Eq. (S32), we have

$$\mu_1^{(D)} - \mu_2^{(D)} = \mu_1^{(A)} - \mu_2^{(A)}. \tag{S33}$$

Substitution of Eq. (S11) into Eq. (S33) yields

$$\frac{x_1^{(D)}}{x_2^{(D)}} = \frac{x_1^{(A)}}{x_2^{(A)}} e^{\frac{\Delta\Delta e}{k_B T}} \tag{S34}$$

or

$$\frac{N_1^{(D)}}{N_2^{(D)}} = \frac{N_1^{(A)}}{N_2^{(A)}} e^{\frac{\Delta\Delta e}{k_B T}}, \tag{S35}$$

where

$$\Delta\Delta e = (a_1 - a_2)\Delta\sigma - (q_1 - q_2)\Delta V. \tag{S36}$$

An important derivation from Eq. (S35) is that the equilibrium lipid distributions are independent of protein distributions in both membranes. This can also be seen from the protein-dependent energy due to the protein-lipid mixing entropy, and according to Eq. (S1), this energy is

$$E_{mixing} = k_B T N_p \ln \frac{N_p}{N_t} \tag{S37}$$

for the membrane containing $N_t$ total lipids and membrane proteins and $N_p$ membrane proteins. Because neither $N_p$ nor $N_t$ changes during the lipid transfer mediated by shuttle LTPs via an lipid exchange mechanism, the energy due to protein-lipid mixing does not change, which leads to the independence of the lipid transfer on the presence membrane proteins in both membranes. In other words, lipid osmosis does not play a role in the equilibrium lipid distributions due to lipid transfer mediated shuttle LTPs.

We derived the equilibrium lipid distributions from Eq.(S35) given experimental control parameters. Let

$$\eta \equiv \frac{N_1^{(D)}}{N_2^{(D)}} = \frac{N_1^{(A)}}{N_2^{(A)}} e^{\frac{\Delta\Delta e}{k_B T}}, \tag{S38}$$

we have

$$\begin{cases} N_1^{(D)} = \eta N_2^{(D)}, \\ N_1^{(A)} = \eta N_2^{(A)} \alpha, \end{cases} \tag{S39}$$

with

$$\alpha \equiv e^{-\frac{\Delta\Delta e}{k_B T}}. \tag{S40}$$

Note that the total number of each species of lipids in both the donor and acceptor membranes $N_1^{(DA)}$ or $N_2^{(DA)}$ is conserved during lipid transfer, i.e.,

$$\begin{cases} N_1^{(D)} + N_1^{(A)} = N_1^{(DA)}, \\ N_2^{(D)} + N_2^{(A)} = N_2^{(DA)}. \end{cases} \tag{S41}$$

Substitution of Eq. (S39) into Eq. (S41) yields

$$\begin{cases} N_1^{(A)} = \dfrac{\left(\eta N_2^{(DA)} - N_1^{(DA)}\right)\alpha}{1-\alpha}, \\ N_2^{(A)} = \dfrac{\eta N_2^{(DA)} - N_1^{(DA)}}{\eta\left(1-\alpha\right)}. \end{cases} \tag{S42}$$

Note that the total number of lipids in each membrane is also conserved during the lipid transfer process, specifically,

$$N_1^{(A)} + N_2^{(A)} = N_L^{(A)}, \tag{S43}$$

where $N_L^{(A)}$ is the total number of lipids in the acceptor membrane. Substitution of Eq. (S42) into Eq. (S43) yields

$$\eta^2 N_2^{(DA)}\alpha + \eta\left[N_2^{(DA)} - N_1^{(DA)}\alpha - \left(1-\alpha\right)N_L^{(A)}\right] - N_1^{(DA)} = 0. \tag{S44}$$

Solving the quadratic equation for $\eta$ and substituting it into Eqs. (S42) and (S41), we can calculate the numbers of lipids in both membranes under equilibrium conditions, given the experimental control parameters $\alpha$, $N_1^{(DA)}$, $N_2^{(DA)}$, and $N_L^{(A)}$.

To facilitate computations, let's again scale the numbers of lipids by the total number of all lipids in two membranes $N_L^{(DA)}$, i.e.,

$$N_L^{(DA)} = N_L^{(D)} + N_L^{(A)} = N_1^{(DA)} + N_2^{(DA)};$$
$$n_i^{(DA)} = \frac{N_i^{(DA)}}{N_L^{(DA)}}; \quad n_L^{(M)} = \frac{N_L^{(M)}}{N_L^{(DA)}}; \quad n_i^{(M)} = \frac{N_i^{(M)}}{N_L^{(DA)}}; \quad M = D, A; \quad i = 1, 2. \tag{S45}$$

Except $n_i^{(M)}$, all other parameters defined here are known experimental control parameters. Then Eq. (S44) can be rewritten as

$$\eta^2 n_2^{(DA)}\alpha + \eta\left[n_2^{(DA)} - n_1^{(DA)}\alpha - (1-\alpha)n_L^{(A)}\right] - n_1^{(DA)} = 0. \tag{S46}$$

In addition, Eq. (S42) can be rewritten as

$$\begin{cases} n_1^{(A)} = \dfrac{\left[\eta - (\eta+1)n_1^{(DA)}\right]\alpha}{1-\alpha}, \\ n_2^{(A)} = \dfrac{\eta - (\eta+1)n_1^{(DA)}}{\eta(1-\alpha)}, \end{cases} \tag{S47}$$

and

$$z_i^{(M)} = \frac{N_i^{(M)}}{N_i^{(DA)}} = \frac{n_i^{(M)}}{n_i^{(DA)}};$$
$$\varsigma_i^{(M)} \equiv \frac{N_i^{(M)}}{N_L^{(M)}} = \frac{n_i^{(M)}}{n_1^{(M)} + n_2^{(M)}}; \quad M = D, A; \quad i = 1, 2. \tag{S48}$$

If the two lipid species have the same lateral areas and charges, or $a_1 = a_2$ and $q_1 = q_2$, then $\Delta\Delta e = 0$ regardless of the differences of membrane tension and potentials between the two membranes. In this case, $\alpha = 1$ and the solution to Eq. (S44) is

$$\eta = \frac{N_1^{(DA)}}{N_2^{(DA)}}. \tag{S49}$$

Substitution of Eq. (S49) into Eqs. (S39) and (S41), we obtained the equilibrium lipid distributions as

$$\frac{N_i^{(D)}}{N_1^{(D)} + N_2^{(D)}} = \frac{N_i^{(A)}}{N_1^{(A)} + N_2^{(A)}} = \frac{N_i^{(DA)}}{N_1^{(DA)} + N_2^{(DA)}}, \quad i = 1, 2. \tag{S50}$$

Therefore, the fraction of each transferable lipid species in both membranes becomes equal and is determined by the fraction of the lipid species in the total transferable lipids in both membranes. In this case, shuttle LTPs catalyze the diffusion of both species of lipids in the two membranes [3].

### 3. Chemical potential of lipids in a vesicular membrane with intrinsic curvature

The membrane of a vesicle of radius $R$ has a bending energy of

$$F = \frac{1}{2}\kappa\left(\frac{2}{R} - \frac{2}{R_s}\right)^2 4\pi R^2 = 8\pi\kappa\left(1 - \frac{R}{R_s}\right)^2, \tag{S51}$$

where $R_s$ is the spontaneous curvature of the membrane and $\kappa$ is its bending rigidity. Suppose the vesicle membrane has zero membrane tension and the lipids in its two leaflets can freely flip, then the total number of lipids in the membrane is

$$N = \frac{8\pi R^2}{a} \tag{S52}$$

where $a$ is the average lateral area per lipid. Expressing the radius $R$ in terms of $N$ and then substituting it into Eq. (S51), we have

$$F = 8\pi\kappa\left(1 - \frac{1}{R_s}\sqrt{\frac{a}{8\pi}}N^{\frac{1}{2}}\right)^2. \tag{S53}$$

The chemical potential of the lipids in the membrane can be calculated as

$$\mu_{bending} = \frac{\partial F}{\partial N} = \frac{2\kappa a}{R_s}\left(\frac{1}{R_s} - \frac{1}{R}\right). \tag{S54}$$

### 4. Membrane curvature generated by lipid osmosis through scramblases

The free energy function of an enclosed lipid bilayer based on the area-difference-elastic (ADE) model is expressed as

$$F = \frac{1}{2}\kappa\oint\left(C_1 + C_2 - C_s\right)^2 dA + \frac{\pi\kappa'}{2h^2 A}\left(\Delta A - \Delta A_s\right)^2, \tag{S55}$$

where $C_1$ and $C_2$ are the principal curvatures of the local membrane curvature, $C_s$ is the spontaneous curvature of the membrane, and $\kappa$ and $\kappa'$ are the local and non-local membrane bending rigidities, respectively [7]. $\Delta A$ and $\Delta A_s$ are the area difference between two leaflets in a stretched and unstretched membrane, respectively, with

$$\Delta A = h\oint\left(C_1 + C_2\right)dA \tag{S56}$$

where $h$ is the distance between the two leaflets.

Let's first calculate the free energy of a vesicle of radius $R$ with no intrinsic membrane curvature ($C_s = 0$) and no area difference between the two relaxed leaflets ($\Delta A_s = 0$). The area difference of the two stretched leaflets in the vesicle in terms of Eq. (S56) is

$$\Delta A = h\frac{2}{R}4\pi R^2 = 8\pi Rh, \tag{S57}$$

and the bending free energy of the vesicle is

$$F = 8\pi\kappa + 8\pi^2\kappa'. \tag{S58}$$

Now let's check how lipid osmosis between two leaflets may generate membrane curvature. Suppose peripheral membrane proteins bind or are lipid-anchored to the outer leaflet of the vesicle with a protein density of $c$. Lipid osmosis causes net lipid flow from the inner leaflet to the outer leaflet, leading to a leaflet tension difference

$$\Delta\sigma = \sigma_- - \sigma_+ = \frac{K\Delta A_{osm}}{2A}, \tag{S59}$$

where $\sigma_-$ and $\sigma_+$ are the membrane tension of the inner leaflet and outer leaflet, respectively, $K$ is the stretching modulus of the lipid bilayer, and $\Delta A_{osm}$ is the area difference between the outer leaflet and the inner leaflet due to lipid osmosis. Here we have assumed that the stretching modulus of a single leaflet is half of that of the lipid bilayer. Under an equilibrium condition, the tension difference equates the osmotic membrane tension, or

$$\Delta\sigma = k_B Tc. \tag{S60}$$

Combining Eqs. (S59) and (S60), we have

$$\frac{\Delta A_{osm}}{A} = \frac{2k_B Tc}{K}. \tag{S61}$$

In the presence of both lipid osmosis and the unstretched area difference, we modify the ADE model in Eq. (S55) as

$$F = \frac{1}{2}\kappa\oint\left(C_1 + C_2 - C_s\right)^2 dA + \frac{\pi\kappa'}{2h^2 A}\left(\Delta A - \Delta A_s - \Delta A_{osm}\right)^2. \tag{S62}$$

To derive the membrane bending induced by lipid osmosis between the two leaflets, we calculate the energy density of a membrane with a mean membrane curvature $C$

$$f \equiv \frac{F}{S} = 2\kappa C^2 + 2\pi\kappa'\left(C - C_m\right)^2 \tag{S63}$$

with

$$C_m \equiv \frac{1}{2h}\left(\frac{\Delta A_s}{A} + \frac{2k_B Tc}{K}\right). \tag{S64}$$

To determine the curvature $C$, we minimize the energy density function in Eq. (S63) with respect to the membrane curvature. To this end, we calculate its derivative

$$\frac{\partial f}{\partial C} = 4\kappa C + 4\pi\kappa'\left(C - C_m\right). \tag{S65}$$

The equilibrium membrane curvature $C_e$ is the solution to the equation

$$\frac{\partial f}{\partial C} = 0, \tag{S66}$$

which yields

$$C_e = \frac{\pi\kappa'}{\kappa + \pi\kappa'}C_m = \frac{\pi\kappa'}{\kappa + \pi\kappa'}\frac{1}{2h}\left(\frac{\Delta A_s}{A} + \frac{2k_B Tc}{K}\right). \tag{S67}$$

Thus, the equilibrium membrane curvature is proportional to both leaflet area difference and the protein density. With $\kappa = \kappa'$, $\Delta A_s = 0$, $c$ = 0.1 nm$^{-2}$, $h$ = 2.5 nm, and $K$ = 250 pN/nm, we have $C_e$ = 5×10$^{-4}$ nm, or the radius of curvature of 2 μm.

### 5. Rate of bulk lipid flow through a model bridge LTP

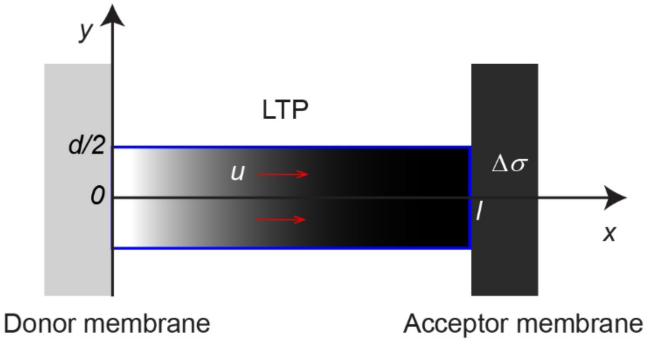


Figure S1. **Model of a bridge LTP.** The right edge of the donor membrane and the left edge of the acceptor membrane are located at $x=0$ and $x=l$, respectively. The acceptor membrane exhibits higher membrane tension ($\Delta\sigma$) and a lower electric potential ($\Delta V$) compared to those of the donor membrane. The bridge LTP is modeled as a lipid conduit with a width $d$ and length $l$. A membrane tension gradient of $\Delta\sigma/l$ drives lipid to flow from the donor membrane to the acceptor membrane.

We model a bridge LTP as a rectangle membrane conduit with a length $l$ and a width $d$ (Figures 1a and S1). The two ends of the conduit are connected to two membranes, one donor membrane and the other acceptor membrane, with a higher membrane tension $\Delta\sigma$ in the acceptor membrane than in the donor membrane. To calculate the lipid velocity profile along the length of the LTP, we choose a Cartesian coordinate such that its origin is located at the center of one end of the LTP and the x-axis points to the other end (Figure S1). In a steady state, the vertical component of the lipid velocity is zero and only the horizontal x component of the lipid flow velocity $u(y)$ is nonzero. In this case, a two-dimensional Navier-Stokes equation can be simplified as

$$\mu\frac{\partial^2 u}{\partial y^2}-\frac{\partial P}{\partial x}+\frac{q\Delta V}{hal}=0, \tag{S68}$$

where $q$ is the average fraction of charge per lipid in the membrane, $a$ is the lateral area of each lipid, and $\mu$ is the viscosity of the lipids in BLTP groove. The pressure $P$ of lipids is the related to membrane tension $\sigma$ by

$$P=-\frac{\sigma}{h}, \tag{S69}$$

where $h$=5 nm is the membrane thickness. We assume that the LTP is long and the first term on the left of Eq. (S68) is independent of $x$, then $\frac{\partial P}{\partial x}$ must be a constant. Based on its boundary condition, we can solve $P$ as

$$\frac{\partial P}{\partial x}=-\frac{\Delta\sigma}{lh} \tag{S70}$$

and rewrite Eq. (S68) as

$$\mu \frac{\partial^2 u}{\partial y^2} + \frac{\Delta\sigma}{lh} + \frac{q\Delta\varphi}{lha} = 0. \tag{S71}$$

The boundary condition of the lipid flow at groove wall of BLTPs is unknown, which depends on interactions between lipids and the wall. We adopt a general Navier partial-slip boundary condition with

$$u\left(\pm\frac{d}{2}\right) = \mp b \left.\frac{\partial u}{\partial y}\right|_{y=\pm\frac{d}{2}}, \tag{S72}$$

where $b$ is the slip length [8]. The slip length is zero for a sticky boundary condition, and infinity for a perfect-slip boundary condition. For water, a slip length up to tens of nanometers was reported for flat and hydrophobic surfaces. Eq. (S68) suggests a quadratic form for its solution, i.e.,

$$u = c_2 y^2 + c_1 y + c_0 \tag{S73}$$

with the three coefficients to be determined. First, according to the symmetry of the system, the velocity profile should be symmetric with respect to x-axis, i.e., $u$ should be an even function with respect to $y$. Thus, $c_1 = 0$. Substituting Eq. (S73) into Eqs. (S71) and (S72), we have

$$c_2 = -\frac{1}{2\mu lh}\left(\Delta\sigma + \frac{q\Delta V}{a}\right) \tag{S74}$$

and

$$c_0 = -\frac{d}{2\mu lh}\left(\Delta\sigma + \frac{q\Delta V}{a}\right)\left(b - \frac{d}{4}\right). \tag{S75}$$

Therefore, we solve the velocity profile as

$$u = \frac{1}{2\mu lh}\left(\Delta\sigma + \frac{q\Delta V}{a}\right)\left(\frac{d^2}{4} - y^2 + bd\right). \tag{S76}$$

The flux of lipid flow can be calculated as

$$\begin{aligned} J &= \frac{2}{a}\int_0^{d/2} u\,dy = -\frac{1}{a\mu lh}\left(\Delta\sigma + \frac{q\Delta V}{a}\right)\int_0^{d/2} y^2 dy + \frac{d}{2a\mu lh}\left(\Delta\sigma + \frac{q\Delta V}{a}\right)\left(\frac{d^2}{4} + bd\right) \\ &= \frac{d^2}{12al\mu h}\left(\Delta\sigma + \frac{q\Delta V}{a}\right)(d + 6b). \end{aligned} \tag{S77}$$

The fluid confined in a nanoscale may exhibit viscosity different from that in bulk [8]. However, in our calculations, we assumed the viscosity of the lipids in BLTP to be the same as that of lipids in the bilayer. The 2D viscosity of lipid bilayers is approximately $\mu h \approx 5.8\times10^{-7}$ pN× s/nm [3]. The slip length of lipids in the hydrophobic groove has not been measured, but is likely small given potentially strong hydrophobic interactions between lipid tails and groove walls [9]. For simplicity, we choose a sticky boundary condition with $b = 0$. The groove width of BLTPs varies from 1 nm to 3 nm along their length. For our estimation, we have chosen an average width of $d$ = 2 nm. Assuming $a = 0.7\,\text{nm}^2$, $l = 20$ nm, and $\Delta\sigma$=0.001-0.1 pN/nm, we estimated the lipid flux as $J$ = 80 – 8208 $s^{-1}$. BLTPs contain charged resides outside the groove region through electroosmosis when a potential difference exists between two membranes [8,10]. This intriguing phenomenon warrants further investigation.

We previously derived the lipid flux through a single bridge LTP assuming a single-file dynamics of lipid flow, which is expressed as

$$J_o = \frac{NaD\Delta\sigma}{k_B T l^2}, \tag{S78}$$

where $N$ is the total number of lipid bound by an LTP and $D$ is the diffusion coefficient of the lipid in the membrane and the LTP [3]. We used the Einstein relation to estimate the 3D viscosity coefficient of the lipid bilayer, or

$$6\pi\mu b = \frac{k_B T}{D}, \tag{S79}$$

where $b$ is the lateral radius of the lipid. Note the number of lipids bound by an LTP (Figure S1) is

$$N = \frac{ld}{a}. \tag{S80}$$

Substituting Eqs. (S79) and (S80) into Eq. (S78), we have

$$J_o = \frac{\Delta\sigma d}{6\pi\mu b l}. \tag{S81}$$

Therefore,

$$\frac{J}{J_o} = \frac{\pi b d^2}{2ah} \approx \frac{d^2}{2bh} \approx 0.84. \tag{S82}$$

Here we have assumed that $a \approx \pi b^2$ with $b = 0.47$ nm.

### 6. Rate of lipid flow through a model bridge LTP estimated from lipid tracer diffusion

To calculate the rate of lipids diffusing through the model LTP shown in Figure S1, we assume that a lipid tracer is present in the donor membrane with an initial number density $\psi_0$ but absent in the acceptor membrane. In addition, the two membranes have equal membrane tension and potential. We derivate the rate of tracer lipid diffusion through the model LTP by adapting the previous derivations [11]. Suppose the tracer density in the channel is $\psi(x)$, then the number of tracer lipids passing the channel position $x$ per unit time and unit length along the y-direction can be written as

$$j = -D\frac{d\psi}{dx}, \qquad [S83]$$

where $D$ is the diffusion constant of the tracer lipid. Because the flux $j$ is a constant and $\psi(0) - \psi(l) = \psi_0$, we can solve for $j$ as

$$j = \frac{D}{l}\psi_0. \qquad [S84]$$

The flux of tracer lipids through the LTP is

$$J_t = jd = \frac{Dd}{l}\psi_0. \tag{S85}$$

An apparent flux of all lipids through the LTP can be calculated by scaling the tracer density to the density of all lipids [12] or replacing $\psi_0$ in Eq. (S85) by $1/a$, yielding

$$J_d = \frac{Dd}{la}. \tag{S86}$$

Note that this apparent flux does not represent any bulk lipid flow. Lipid diffusion can occur without any bulk lipid flow or membrane area changes, as illustrated by Figure 3d. Although lipid diffusion causes the net transfer of a specific species of lipids such as the tracer lipids from one membrane to another membrane, other lipid species may transfer in an opposite direction to counteract bulk lipid flow.

### 7. Supplement Text References:


1. Phillips R, Kondev J, Theriot J, Garcia H: *Physical Biology of the Cell* edn 2nd: Garland Science; 2012.
2. Jahnig F: **Lipid exchange between membranes**. *Biophysical Journal* 1984, **46**:687-694.
3. Zhang YL, Ge J, Bian X, Kumar A: **Quantitative models of lipid transfer and membrane contact formation**. *Contact* 2022, **5**:1-21.
4. Chung J, Torta F, Masai K, Lucast L, Czapla H, Tanner LB, Narayanaswamy P, Wenk MR, Nakatsu F, De Camilli P: **PI4P/phosphatidylserine countertransport at ORP5- and ORP8-mediated ER-plasma membrane contacts**. *Science* 2015, **349**:428-432.
5. de Saint-Jean M, Delfosse V, Douguet D, Chicanne G, Payrastre B, Bourguet W, Antonny B, Drin G: **Osh4p exchanges sterols for phosphatidylinositol 4-phosphate between lipid bilayers**. *Journal of Cell Biology* 2011, **195**:965-978.
6. Wong LH, Gatta AT, Levine TP: **Lipid transfer proteins: the lipid commute via shuttles, bridges and tubes**. *Nature Reviews Molecular Cell Biology* 2019, **20**:85-101.
7. Seifert U: **The concept of effective tension for fluctuating vesicles**. *Zeitschrift Fur Physik B-Condensed Matter* 1995, **97**:299-309.
8. Kavokine N, Netz RR, Bocquet L: **Fluids at the Nanoscale: From Continuum to Subcontinuum Transport**. *Annual Review of Fluid Mechanics* 2021, **53**:377-410.
9. Liu C, Li ZG: **On the validity of the Navier-Stokes equations for nanoscale liquid flows: The role of channel size**. *Aip Advances* 2011, **1**.
10. Bocquet L, Charlaix E: **Nanofluidics, from bulk to interfaces**. *Chemical Society Reviews* 2010, **39**:1073-1095.
11. Kargol A: **Ion Channels**. In *Introduction to Cellular Biophysics, Volume 1: Membrane Transport Mechanisms*. Edited by: Iop Concise Physics; 2019.
12. Ohmann A, Li CY, Maffeo C, Al Nahas K, Baumann KN, Gopfrich K, Yoo J, Keyser UF, Aksimentiev A: **A synthetic enzyme built from DNA flips $10^7$ lipids per second in biological membranes**. *Nature Communications* 2018, **9**:2426.
